# Ectopic Expression of DNA Repair Enzymes Modulates Survival following Ultraviolet Irradiation Challenge

**DOI:** 10.1101/185058

**Authors:** Stuti P. Garg, Irina G. Minko, Erdem Coskun, Onur Erdem, Pawel Jaruga, Miral Dizdaroglu, R. Stephen Lloyd

## Abstract

In *Escherichia coli*, the nucleotide excision repair (NER) pathway removes ultraviolet (UV) light-induced cyclobutane pyrimidine dimers (CPDs) and 6-4 dipyrimidine photoproducts (6-4 PPs). Activation of alternative repair pathways, such as base excision repair (BER) and nucleotide incision repair (NIR), is inoperative because this organism lacks both the necessary BER DNA glycosylase and NIR UV endonuclease to initiate repair of these lesions. To determine if initiation of either pathway would enhance survival to biologically-relevant UV irradiation, the BER and NIR pathways were activated by expression of *Chlorella* virus-1 pyrimidine dimer glycosylase (cv-pdg) and *Schizosaccharomyces pombe* UV endonuclease (UVDE), respectively. The substrate specificity of cv-pdg includes CPDs and ring-fragmented purines, 4,6-diamino-5-formamidopyrimidine and 2,6-diamino-4-hydroxy-5-formamidopyrimidine, but not 6-4 PPs. In contrast, while UVDE incises DNA containing CPDs and 6-4 PPs, it was not previously known if the substrate specificity of UVDE included DNA containing ring-fragmented purines. Mass spectrometry was used to establish that these oxidatively-induced lesions were not substrates for UVDE. Expression of either cv-pdg or UVDE in NER-deficient *E. coli* significantly enhanced survival following UVB irradiation, but not to the levels of wild type (WT) cells. Survival of NER-proficient, homologous recombination-deficient cells could also be significantly enhanced by expression of either enzyme, suggesting that in response to UVB exposure, interactions between NER and activated BER or NIR pathways could be additive. Further, expression of cv-pdg or UVDE in WT *E. coli* enhanced survival following solar-simulated light (SSL) exposures.

## 1. Introduction

The spectrum of ultraviolet (UV) irradiation can be divided into three major regions based on differences in photochemistry and biological interactions. UVC irradiation (<280 nm) is strongly absorbed by DNA, producing cyclobutane pyrimidine dimers (CPDs), 6-4 photoproducts (6-4 PPs), and many other DNA lesions. However, these wavelengths are not germane to UV exposures at the Earth’s surface due to atmospheric attenuation. UVB (280-315 nm) is the most biologically damaging component of sunlight that reaches the Earth’s surface. Although it accounts for <5% of the UV that passes through the ozone, the UVB causes direct photochemical DNA damage, including CPDs and 6-4 PPs [1]. The role of these lesions in UV-induced mutagenesis has been established, particularly in human clinical samples of skin tumors and pre-cancerous lesions [2]. In addition, UVB has been demonstrated to oxidatively induce DNA damage, both directly and via reactive oxygen species [3,4]. Pure DNA only weakly absorbs UVA (315-400 nm), but in cells, endogenous photosensitizers amplify the DNA damaging effects of these wavelengths, resulting in the formation of the same DNA lesions induced following exposure to shorter wavelengths [1,4,5]. UVA has been implicated in photoaging, immunosuppression, and the generation of reactive oxygen species [3,6-8].

In *Escherichia coli (E. coli)*, DNA repair of CPDs and 6-4 PPs primarily proceeds via the Nucleotide Excision Repair (NER) pathway, with the initiation of this pathway requiring the coordinated activities of the UvrA, UvrB, and UvrC proteins. Defects in any of these proteins confer significant increases in UV-induced cytotoxicity [9,10]. Additionally, CPDs can undergo photoreversal under appropriate conditions through the activity of photolyase [10]. The homologous recombination (HR) and translesion DNA synthesis pathways that constitute damage tolerance mechanisms rather than repair processes, are also important for cellular survival following UV exposure [11].

While not present in *E. coli*, many bacteria and viruses utilize DNA glycosylases to initiate repair of CPDs via the Base Excision Repair (BER) pathway [12,13]. These enzymes cleave the *N*-glycosidic bond of the 5’ pyrimidine of CPD and subsequently incise the phosphodiester backbone at the abasic site via a lyase mechanism [14-16]. Several bacteriophage species of the *Myoviridae* and *Phycodnaviridae* (*Chlorella* virus) families possess such pyrimidine dimer glycosylases (pdgs) [17-20], and similar sequences have been found in bacterial genomes, only distantly related to each other (*Brucella, Prochlorococcus, Bordetella, Haemophilus, Pasteurella*) [21-23]. The best characterized of the viral enzymes are the T4 and *Paramecium bursari chlorella* virus-1 pdgs (T4-pdg and cv-pdg, respectively) [24-27]. In addition to CPDs, cv-pdg has been reported to catalyze incision of a subset of oxidatively-induced DNA lesions, including 4,6-diamino-5-formamidopyrimidine (FapyAde) and 2,6-diamino-4-hydroxy-5-formamidopyrimidine (FapyGua) [26]. The formation of these ring-fragmented purines in UVB- and UVC-irradiated DNA in aqueous solution and in the skin of UVB-irradiated mice has been demonstrated [28,29]. Since *E. coli* has all necessary BER components to complete glycosylase-initiated repair, it was hypothesized that complementation with pdgs would enhance survival following UV exposure. Indeed, partial complementation has been demonstrated for UVC exposure of an *E. coli* strain that was defective in both NER and HR [30-32].

In addition to the use of BER to augment or substitute for *E. coli* NER, the Nucleotide Incision Repair (NIR) pathway for UV-induced photoproducts can be initiated by the activity of a UV damage-specific endonuclease (UVDE) [33]. *Schizosaccharomyces pombe* UVDE has a broad substrate range that includes CPDs, 6-4 PPs, abasic sites, small loops, and mismatches [34-36]. However, substrate specificity of UVDE regarding oxidatively-induced DNA base damage has not been previously characterized, except for the demonstration of a weak activity on DNA containing a site-specific dihydrouracil paired with non-cognate Gua as opposed to Ade [34]. In contrast to the glycosylase mechanism of pdgs, UVDE is an endonuclease that cleaves the DNA phosphodiester backbone 5’ to the damage site, leaving a 5’ phosphate and a 3’ OH, thus initiating the NIR pathway.

Although prior biochemical and structural biology investigations have extensively characterized the pdg and UVDE enzymes [24-27,34-36], the biological efficacies of activating either pathway in response to exposures to biologically relevant UV wavelengths have not been performed. Thus, the abilities of cv-pdg and UVDE to alter biological endpoints associated with UVB and solar simulated light (SSL) exposures were examined following expression of these enzymes in *E. coli* strains that either were wild type (WT) for DNA repair and tolerance pathways or had a deficiency in NER or HR.

## 2. Materials and Methods

### 2.1. UVB and SSL sources

UVB irradiation was performed using an Ultraviolet Products 302 nm UV lamp (15-watt) without filter. The intensity was determined using an IL1400A meter (International Light INC) with the SEL204/UVB/W detection probe (measurement range: 275-310 nm) and the doses were calculated. SSL was produced by a Newport Class ABA 1600 W Solar Simulator, with atmospheric attenuation filter. The intensities of UVA and UVB components were separately measured using SEL033/UVA/TD (measurement range: 315-390 nm) and SEL204/UVB/W detection probes. The dose rates for UVA and UVB were ~5.7 and 0.2 kJ/m^2^/min, respectively, for the fixed-height irradiation. The relative yield of UVA and UVB were ~96 and 4%, respectively. All exposures to SSL were performed under these standard conditions. For simplicity, the data are reported as a function of irradiation time.

### 2.2. E. coli strains

The genotypes of the *E. coli* strains used in the UVB and SSL analyses are given in Table 1. The parental AB1157 strain and isogenic mutants that were either defective in NER (AB2500, *uvrA6*) or HR (AB2487, *recA* 13) were obtained from the *E. coli* Genetic Stock Center (Yale University). The DH5a and BL21 (DE3) *E. coli* strains used for cloning and initial characterization of DNA constructs were purchased from Invitrogen.

**Table 1.** *E. coli* strains used in this study

| **Name** | **Genotype** |
| --- | --- |
| **AB2500** | F<sup>-</sup>, *thr-1*, *araC14*, *leuB6*(Am), $\Delta$(*gpt-proA*)62, *lacY1*, *tsx-33*, *qsr'-0*, *glnX44*(AS), *galK2*(Oc), $\lambda^-$, *hisG4*(Oc), *thyA15*, *rpsL31*(strR), *xylA5*, *mtl-1*, *argE3*(Oc), *thiE1*, ***uvrA6***, *deoB12* |
| **AB1157** | F<sup>-</sup>, *thr-1*, *araC14*, *leuB6*(Am), $\Delta$(*gpt-proA*)62, *lacY1*, *tsx-33*, *qsr'-0*, *glnX44*(AS), *galK2*(Oc), $\lambda^-$, *Rac-0*, *hisG4*(Oc), *rfbC1*, *mgl-51*, *rpoS396*(Am), *rpsL31*(strR), *kdgK51*, *xylA5*, *mtl-1*, *argE3*(Oc), *thiE1* |
| **AB2487** | F<sup>-</sup>, *thr-1*, *araC14*, *leuB6*(Am), $\Delta$(*gpt-proA*)62, *lacY1*, *tsx-33*, *qsr'-0*, *glnX44*(AS), *galK2*(Oc), $\lambda^-$, *Rac-0*, *hisG4*(Oc), *rfbC1*, ***recA13***, *thyA16*, *rpsL31*(strR), *kdgK51*, *xylA5*, *mtl-1*, *argE3*(Oc), *thiE1*, *deoB11* |

### 2.3. Cell survival following exposure to UVB or SSL

Following overnight growth in LB media supplemented with ampicillin (100 µg/mL), cells were serially diluted. Equal quantities of multiple dilutions were aliquoted in triplicate onto prewarmed LB-ampicillin plates and the media allowed to absorb into the agar. Following irradiation with either UVB or SSL, plates were incubated at 30 °C for 24-48 h, until colonies were ~2 mm in diameter. Only plates containing >25 colonies were analyzed. All experiments were repeated a minimum of 3 times after establishing the optimal exposure conditions for each strain and source of irradiation. Data are expressed as means with standard errors. The *p* values were calculated using two-tailed Student’s t-test. Values are considered significant for *p* ≤ 0.05.

### 2.4. Cloning and expression of cv-pdg and UVDE

The expression construct for cv-pdg was previously described [37]. The gene encoding the catalytically-active, N-terminally truncated form of UVDE (Δ228) (gift from Dr. Paul Doetsch, Emory University) was inserted into a pET-22b vector using unique *Nde* I and *Hind* III sites, and the sequence of the cloned gene was verified by Sanger sequencing. Plasmid expression vectors were introduced into BL21 (DE3) cells per the supplier’s protocol, and crude cellular lysates were prepared from the overnight cultures to assay for steady-state expression of active enzymes. Expression of the enzymes was confirmed by incubation of cellular lysates with UVC-irradiated plasmid DNA (data not shown). The pET-22b vectors containing the *cv-pdg* or *uvde(Δ228)* gene or no insert were introduced by electroporation into AB1157, AB2500, and AB2487 cells. Ampicillin resistant clones were selected and stored as frozen glycerol stocks. Cultures used for cell survival analyses were made from fresh single-colony isolates derived from glycerol stocks.

### 2.5. Preparation of DNA samples

Commercially available calf thymus DNA was dissolved in phosphate buffer (pH 7.4) (0.3 mg/mL) at 4 ºC. The DNA solution was saturated with N_2_O for 30 min, irradiated with γ-rays in a ^60^Co γ ray-source at a dose of 10 Gy (dose rate 5.17 Gy/min), and DNA was dialyzed against water at 4 °C for 18 h. After dialysis, the DNA concentration was measured by absorption spectrophotometry. Aliquots (50 μg) of the DNA sample were supplemented with aliquots of the internal standards, i.e., FapyAde-^13^C,^15^N_2_, FapyGua-^13^C,^15^N_2_, 8-OH-Ade-^13^C,^15^N_2_, 8-OH-Gua-^15^N_5_, 5-OH-Cyt-^13^C,^15^N_2_, 5-OH-Ura-^13^C_4_,^15^N_2_, ThyGly-^2^H_4_, 5-OH-5-MeHyd-^13^C,^15^N_2_ and 5,6-diOH-Ura-^13^C,^15^N_2_ (isodialuric acid-^13^C,^15^N_2_), which are a part of the NIST Standard Reference Material 2396 Oxidative DNA Damage Mass Spectrometry Standards (for details see http://www.nist.gov/srm/index.cfm and https://www-s.nist.gov/srmors/view_detail.cfm?srm=2396). The samples were dried in a SpeedVac under vacuum and kept at 4 °C until use.

For the enzymatic treatment of DNA samples, two different buffer solutions were used: Buffer 1 (no divalent metal ions): 50 mM phosphate buffer (pH 7.4), 100 mM KCl, 1 mM EDTA, and 0.1 mM dithiothreitol; Buffer 2 (plus divalent metal ions): 50 mM phosphate buffer (pH 7.4), 100 mM KCl, 2 mM MnCl_2_, 2 mM MgCl_2_, 1 mM EDTA, and 0.1 mM dithiothreitol.

DNA samples (50 μg) were dissolved in 50 μL of an incubation buffer and incubated with 5 μg of UVDE or 5 μg cv-pdg for 1 h at 37 °C. Another set of samples was sequentially incubated with 5 μg of UVDE for 1 h and with 5 μg of cv-pdg for an additional 1 h. DNA samples to be used as controls were incubated without any enzyme. Four independently prepared samples were used for each data point. After incubation, 125 μL of cold ethanol (kept at −20 °C) were added to the samples to stop the reaction and to precipitate DNA. The samples were kept at −20 °C for 1 h, and then centrifuged at 15,000 g for 30 min at 4 °C. DNA pellets and supernatant fractions were separated. Ethanol was removed from the supernatant fractions under vacuum in a SpeedVac. Aqueous supernatant fractions were frozen at −80 °C for 1 h and then lyophilized to dryness for 18 h.

### 2.6. Measurement of DNA base lesions by gas chromatography-tandem mass spectrometry (GC-MS/MS)

To each lyophilized supernatant fraction, 60 μL of a mixture of nitrogen-bubbled bis(trimethylsilyl)trifluoroacetic acid [containing trimethylchlorosilane (1%; v/v)] (BSTFA) (Pierce, Rockford, Ill.) and pyridine (Sigma-Aldrich, St. Louis, MO) (1:1, v/v) was added. The samples were vortexed and purged individually with ultra-high-purity nitrogen, tightly sealed with Teflon-coated septa, and then heated at 120 °C for 30 min. After cooling, the clear samples were removed and placed in vials used for injection onto the GC-column. Vials were purged with ultra-high-purity nitrogen and tightly sealed with Teflon-coated septa. Aliquots (4 μL) of derivatized samples were analyzed by GC-MS/MS under the conditions described previously [38]. On the basis of the known mass spectra of trimethylsilyl derivatives of DNA base lesions [39-41], the following mass transitions were used for identification and quantification: *m/z* 369 → *m/z* 280 and *m/z* 372 → *m/z* 283 for FapyAde and FapyAde-^13^C,^15^N_2_ respectively; *m/z* 457 → *m/z* 368 and *m/z* 460 → *m/z* 371 for FapyGua and FapyGua-^13^C,^15^N_2_, respectively; *m/z* 367 → *m/z* 352 and *m/z* 370 → *m/z* 355 for 8-OH-Ade and 8-OH-Ade-^13^C,^15^N_2_, respectively; *m/z* 455 → *m/z* 440 and *m/z* 460 → *m/z* 445 for 8-OH-Gua and 8-OH-Gua-^15^N_5_, respectively; *m/z* 343 → *m/z* 342, and *m/z* 346 → *m/z* 345 for 5-OH-Cyt and 5-OH-Cyt-^13^C,^15^N_2_, respectively; *m/z* 344 → *m/z* 343, and *m/z* 350 → *m/z* 349 for 5-OH-Ura and 5-OH-Ura-^13^C_4_,^15^N_2_, respectively; *m/z* 448 → *m/z* 259 and *m/z* 452 → *m/z* 262 for ThyGly and ThyGly-^2^H_4_, respectively; *m/z* 331 → *m/z* 316 and *m/z* 334 → *m/z* 319 for 5-OH-5-MeHyd and 5-OH-5-MeHyd-^13^C,^15^N_2_, respectively; *m/z* 432 → *m/z* 417 and *m/z* 435 → *m/z* 420 for 5,6-diOH-Ura and 5,6-diOH-Ura-^13^C,^15^N_2_, respectively. The quantification was achieved using integrated areas of the signals of the mass transitions of the monitored DNA base lesions and those of their stable isotope-labeled analogues.

## 3. Results and Discussion

### 3.1. Experimental Rationale and Design

DNA repair of CPDs and 6-4 PPs in *E. coli* is limited to NER, except for CPD photoreversal under appropriate activation conditions. However, analyses of DNA repair pathways in certain viruses and other prokaryotic organisms reveal that alternative repair mechanisms, including BER and NIR are possible, but in *E. coli*, key enzymes are not present to initiate either pathway for the repair of UV-induced dipyrimidine photoproducts. To gain an understanding of how these alternative pathways may modulate cell survival in DNA repair-proficient and -deficient *E. coli*, we implemented a strategy to activate the BER pathway for CPDs and the ring-fragmented purines, FapyAde and FapyGua, using cv-pdg. In parallel, NIR was activated for CPDs and 6-4 PPs by expression of S. *pombe* UVDE. Previous investigations have consistently revealed that expression of T4-pdg [30-32] or cv-pdg [24] in UVC-irradiated *E. coli* AB2480 (both NER- and HR-deficient), resulted in increased survival to levels approaching that observed with either the NER or HR single mutant, but not to WT levels. However, consequences of activating BER pathway using biologically relevant wavelengths of UV light have not been studied. The alteration of cellular responses via activation of the NIR pathway in *E. coli* has not been addressed under any conditions of UV exposure. Such analyses have the potential to distinguish the relative importance of repair of oxidatively-induced DNA damage versus 6-4 PP repair in a bacterial context.

### 3.2. FapyGua and FapyAde are not substrates for UVDE

UVDE has been previously shown to incise DNAs containing CPDs, 6-4 PPs, abasic sites, small loops, mismatches and other DNA lesions [34-36]. To understand the contribution of UVDE-initiated repair to survival following UVB and SSL exposures, we examined the ability of UVDE to release FapyGua or FapyAde from γ-irradiated DNA. Since we had previously demonstrated using GC-MS methods, that cv-pdg efficiently excised these ring-fragmented purines [26], we exploited these activities to determine whether UVDE could incise DNA at these sites. To serve as a control for release of FapyGua or FapyAde, purified cv-pdg was incubated with γ-irradiated calf thymus DNA in buffers containing either no metal ions or MgCl_2_ and MnCl_2_. As anticipated, cv-pdg released FapyAde and FapyGua (Fig. 1, Panels A and B, respectively) in the absence or presence of MgCl_2_ and MnCl_2_. Identical reactions were also performed using UVDE that exhibits maximum activity in the presence of these metal ions. Since UVDE has been previously characterized as an endonuclease, not a glycosylase, and this assay measures free base release, it was not surprising that levels of FapyAde and FapyGua were indistinguishable from the no enzyme controls following incubation in either the presence or absence of the divalent metal ions (Fig. 1, Panels A and B). To determine if UVDE had incised the DNA at fragmented purines, DNA was incubated with UVDE for 1 h prior to the addition of cv-pdg. It was reasoned that if UVDE incised DNA at sites containing FapyAde or FapyGua, these would not be further cleavable as result of subsequent addition of cv-pdg. However, if FapyAde and FapyGua remained intact, subsequent reaction with cv-pdg should release equivalent amounts of these base adducts as compared to incubation with cv-pdg alone. Data generated from the sequential addition of UVDE for 1 h and cv-pdg for an additional 1 h (UVDE + cv-pdg) showed that the same quantity of both lesions was released, demonstrating that UVDE does not incise DNA at either FapyAde or FapyGua sites (Fig. 1, Panels A and B, respectively). Thus, although cv-pdg and UVDE can both initiate repair of CPDs, these enzymes have distinct substrate specificities. While UVDE incises DNA containing 6-4 PPs, cv-pdg catalyzes removal of FapyAde and FapyGua.

**Figure 1.**
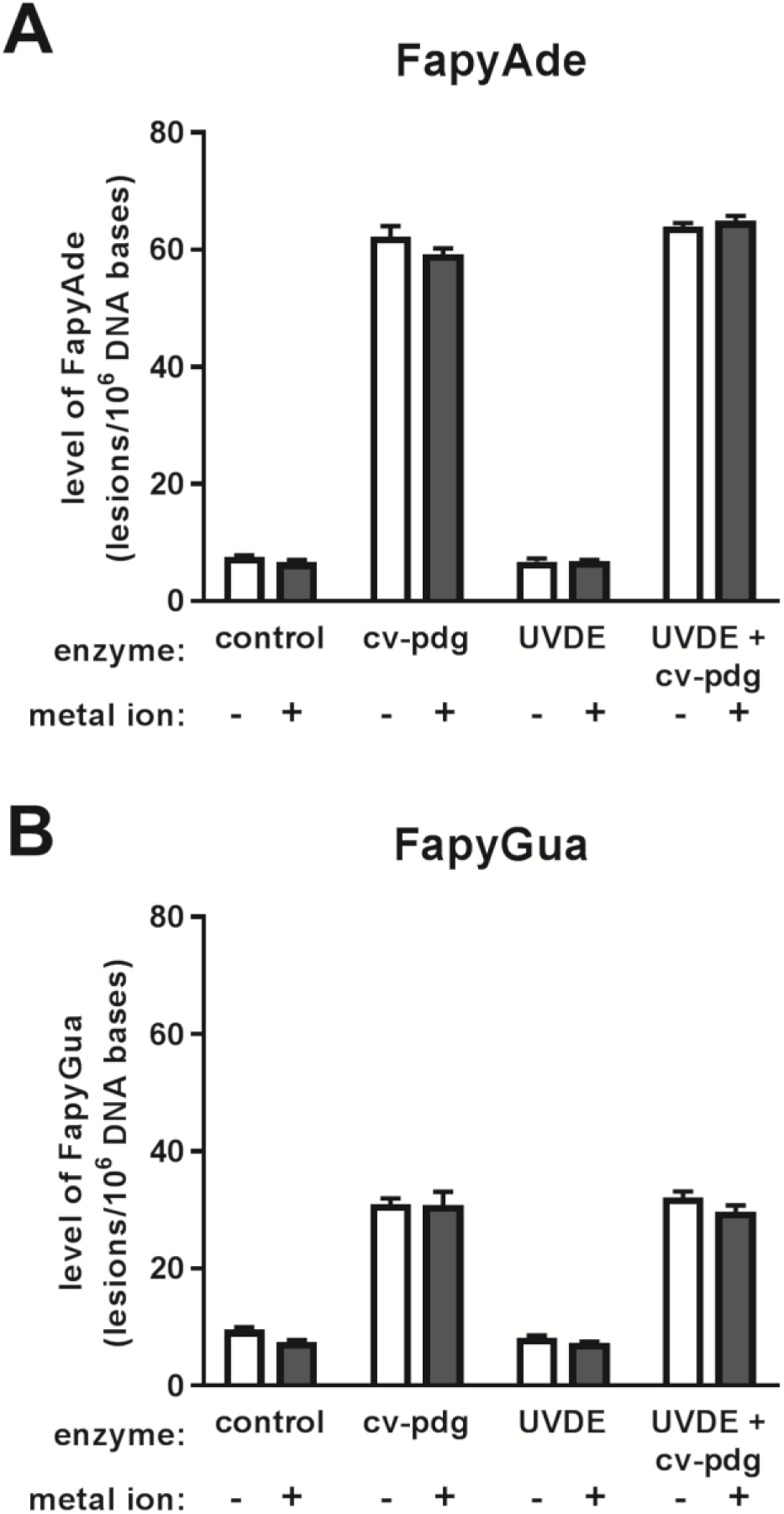
Lack of excision of FapyAde and FapyGua by UVDE from y-irradiated DNA. Calf thymus DNA in a N2O-saturated phosphate buffer was irradiated with a 10 Gy dose of y-rays using a ^60^Co y ray-source, dialyzed, and 50 μg aliquots incubated with cv-pdg alone, UVDE alone, or sequential addition of UVDE for 1 h followed by cv-pdg. The amounts of FapyAde (Panel A) and FapyGua (Panel B) released were measured by GC-MS/MS. The incision reactions were performed in the absence or presence of MnCl_2_ and MgCl_2_. Four independently prepared samples were used for each data point. The uncertainties are standard deviations.

### 3.3. Modulation of E. coli survival following UVB exposure by expression of cv-pdg or UVDE

Having established the substrate specificities of cv-pdg and UVDE, we determined if expression of these enzymes in repair-proficient or -deficient *E. coli* could modulate cell survival following exposure to UVB. Plasmids were constructed that positioned the genes encoding cv-pdg and UVDE downstream of a T7-RNA polymerase promoter that produces low, steady-state levels of each of these enzymes. These un-induced levels of expression of cv-pdg and UVDE were sufficient to exert a robust biological response in NER-deficient *E. coli* cells following challenge to irradiation as determined by UVB (and SSL) gradient dose responses, but did not cause noticeable toxicity under normal growth conditions (data not shown).

To quantify the effect of cv-pdg and UVDE on survival of *E. coli* following UVB irradiation, serial dilutions of WT and NER-deficient (*UvrA*^-^) cells were challenged with increasing doses of UVB and colony formation was measured (Fig. 2, Panels A and B, respectively). As expected, the NER-deficient strain was much more sensitive than WT to UVB exposure, with equivalent doses required to reduce survival to 37% being ~8 and 79 J/m^2^, respectively. Expression of either cv-pdg or UVDE did not enhance survival of WT *E. coli* at any of the doses examined (Panel A). In contrast, both cv-pdg and UVDE significantly improved survival of the NER-deficient cells at all doses (*p* = 0.00000062 and 0.00062 at 10 J/m^2^, 0.0000014 and 0.0000019 at 20 J/m^2^, and 0.00038 and 0.00020 at 30 J/m^2^, respectively). Enhancement conferred by cv-pdg was greater than that measured for ectopic expression of UVDE (*p* = 0.010 at 10 J/m^2^, 0.00013 at 20 J/m^2^, and 0.0033 at 30 J/m^2^). Since this phenomenon was observed at all three doses tested, it is unlikely to be explained by possible differences in either the expression levels of active enzymes or availabilities of downstream activities to complete BER or NIR. An alternative explanation could be that there may only be a minor contribution of 6-4 PPs to the cytotoxic effects of UVB in *E. coli* and that enhanced repair of ring-fragmented purines by cv-pdg may be more important for tolerating UVB. This assumption would be consistent with the observations that in both isolated and cellular DNAs exposed to UVB, 6-4 PPs are generated at a much lower frequency than CPDs (reviewed in [1]). Ectopic expression of either enzyme did not fully restore survival of the UvrA-deficient cells to that of WT, with equivalent doses required to reduce survival to 37% being 16 and 28 J/m^2^ in UvrA-deficient cells expressing UVDE and cv-pdg respectively, relative to 79 J/m^2^ in WT. These data suggest that the initiation of alternative pathways such as BER and NIR, could not fully substitute for the intact NER system of *E. coli*.

**Figure 2.**
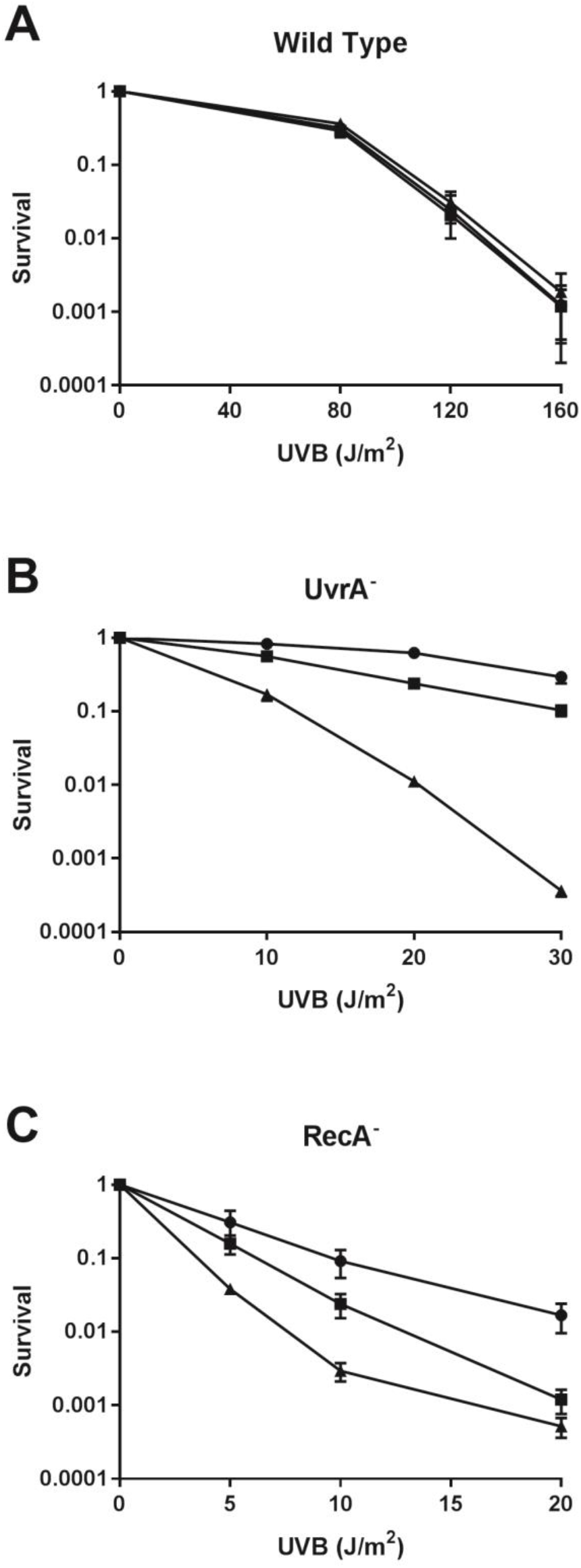
Survival of WT and repair-deficient *E. coli* strains that express enzymes to activate BER and NIR pathways for UVB-induced DNA damage. WT (Panel A), UvrA-deficient (Panel B), and RecA-deficient (Panel C) *E. coli* that expressed cv-pdg (circles) or UVDE (squares) or contained an empty vector (triangles), were irradiated with increasing levels of UVB and colony forming ability measured. At least three independent experiments were used for each data point. The data are presented as means with standard errors. The *p* values were calculated using two-tailed Student’s t-test.

The observation that there was no enhanced survival following UVB exposure in WT cells containing plasmids for the expression of cv-pdg or UVDE, raised the concern that NER may compete with BER and NIR for either common DNA substrates, shared downstream activities, or other factors that limit repair. Thus, experiments were designed to examine whether it was possible to increase survival by expressing these enzymes in any other NER-proficient strain. Since HR-deficient cells are also known to be UVB hypersensitive [42,43], *E. coli* lacking functional RecA, an essential HR component [11], was used to address this question. RecA-deficient cells were transformed with the same set of three plasmids and following selection, were challenged with increasing doses of UVB (Fig. 2, Panel C). These data showed that even though the NER pathway was intact, activation of either cv-pdg- or UVDE-initiated repair enhanced survival (*p* = 0.11 and 0.067 at 5 J/m^2^, 0.031 and 0.0085 at 10 J/m^2^, and 0.036 and 061 at 20 J/m^2^, respectively). Also, consistently observed at all doses, but not statistically significant, there was a trend toward greater enhancements in survival in cells expressing cv-pdg versus UVDE. These data reveal that in the absence of HR, survival of *E. coli* as conferred by NER can be augmented by activation of either BER or NIR.

### 3.4. Modulation of E. coli survival following SSL exposure by expression of cv-pdg or UVDE

While the investigation described above focused on the biological effects of cellular response to a very narrow range of UVB wavelengths, terrestrial exposure to UV irradiation is significantly more complex and can be modelled by SSL. Our SSL experimental system generates ~4% UVB relative to the total of UVA and UVB. To determine if expression of either cv-pdg or UVDE would modulate survival of WT following SSL exposure, cultures were exposed to increasing durations of SSL and the colony-forming abilities were measured (Fig. 3). Although survival of WT *E. coli* expressing UVDE was not significantly different from control at the 8-min and 16-min exposures, it was significantly enhanced at the 24-min exposure (*p* = 0.034). The effect of expression of cv-pdg was even greater, with significant differences observed at the 16-min and 24-min exposures (*p* = 0.025 and 0.020, respectively) (Fig. 3). Similar analyses were carried out in UvrA-deficient cells, with the trends of enhanced survival observed for cells complemented with either cv-pdg or UVDE at the 24 min exposure (data not shown). Thus, the spectra of SSL-induced DNA damage include a fraction of cytotoxic lesions that can be repaired via initiation of BER or NIR. Since enhancement of survival by expression of cv-pdg or UVDE was observed in WT cells (Fig. 3), the cytotoxic SSL-induced lesions that represent substrates for these repair enzymes, are inefficiently, if not at all, recognized by NER. This conclusion agrees with prior studies demonstrating that the major component of solar UV, UVA, causes oxidative stress conditions [4-6] and that many common oxidatively-induced DNA lesions represent poor substrates for NER [9].

**Figure 3.**
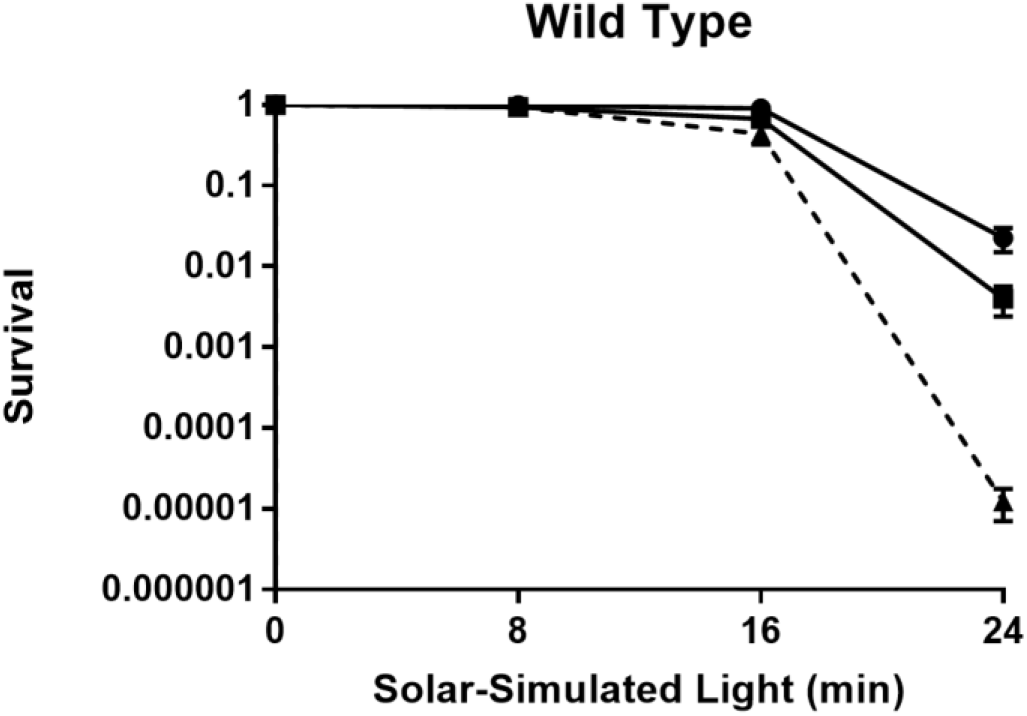
Survival of WT *E. coli* that express enzymes to activate BER and NIR pathways for SSL-induced DNA damage. WT *E. coli* that expressed cv-pdg (circles) or UVDE (squares) or contained an empty vector (triangles), were irradiated with increasing levels of SSL and colony forming ability measured. At least three independent experiments were used for each data point. The data are presented as means with standard errors. The *p* values were calculated using two-tailed Student’s *t*-test.

## 4 Conclusions

The data presented herein demonstrate the efficacy of enhancing the DNA repair capacity in WT and repair-deficient *E. coli* cells following exposures to UVB irradiation and to a limited extent, SSL. Extrapolation of these prokaryotic data suggest that mammalian cells might also benefit from enhanced DNA repair capacity of CPDs, 6-4 PPs, or oxidatively-induced DNA damage caused by UV exposure.

## Acknowledgements

We wish to thank Dr. Paul W. Doetsch, Emory University for the expression plasmid for the truncated form of UVDE, Drs. Vladimir Vartanian, Nichole Owen, and Anuradha Kumari for laboratory and figure presentation assistance, and Dr. Amanda McCullough for guidance and helpful consultations throughout these investigations. Certain commercial equipment or materials are identified in this paper to specify adequately the experimental procedure. Such identification does not imply recommendation or endorsement by the National Institute of Standards and Technology, nor does it imply that the materials or equipment identified are necessarily the best available for the purpose.

